# Breaking barriers: A new phytomedicine based treatment approach for targeted unrevealed non-canonical DNA structures in tuberculosis bacteria

**DOI:** 10.1101/2025.04.08.647720

**Authors:** Sudipta Bhowmik, Tania Shroff, Sagar Bag, Pratibha Kadam, Nerges Mistry

## Abstract

Tuberculosis (TB), caused by *Mycobacterium tuberculosis* (*Mtb*), remains a critical global health issue, complicated by the emergence of multi-drug-resistant (MDR) and extensively drug-resistant (XDR) strains. Current treatments involve prolonged use of first- and second-line drugs, which are associated with severe side effects and poor patient adherence. Non-canonical DNA structures, such as G-quadruplexes (GQ) and i-Motifs (iM), play a vital role in regulating *Mtb’s* virulence, stress responses, and drug resistance mechanisms, making them attractive targets for therapy. Flavonoids, naturally occurring polyphenolic compounds found in various fruits and vegetables have demonstrated the ability to enhance the effectiveness of traditional TB drugs while minimizing cytotoxicity.

By employing biophysical techniques including UV-Vis absorption spectroscopy, binding constant determination, and thermal melting experiments, this study investigates the interaction between two flavonoids, quercetin and kaempferol and non-canonical DNA structures (GQ and iM) within the *Mtb* genome. Key GQ/iM sequences from *Mtb* genes associated with drug resistance were identified and evaluated for their binding with flavonoids. Results revealed that quercetin and kaempferol preferentially interact with specific cyp51 GQ, dnaB GQ, espB GQ, espE GQ, SigA iM, fabH iM, psk5 iM DNA sequences, as indicated by significant changes in absorption spectra. The calculated binding constants showed strong affinities for these specific DNA structures. Thermal melting experiments further indicated that flavonoids increased the thermal stability of these particular GQ and iM DNA, suggesting stabilization via strong stacking interactions. These findings highlight the potential of flavonoids as promising agents to target GQ/iM DNA structures, offering a new strategy for addressing *Mtb* drug resistance and virulence.

## Introduction

Tuberculosis (TB) caused by *Mycobacterium tuberculosis* (*Mtb*) remains a major global health challenge, with rising multi-drug-resistant (MDR) and extensively drug-resistant (XDR) strain [1]. The rise of MDR or XDR *Mtb* poses a significant threat to the treatment efficacy of TB therapies. Current therapies target mainly essential cellular processes by applying a combination of first-& second-line anti-TB drugs taken over 9-18 months [2]. This long duration is accompanied by frequent severe side effects which leads to unsatisfactory patient compliance and withdrawal from treatment further contributing to resistance[2]. Identifying and targeting key genes and sequences associated with survival and resistance, aids in the development of targeted therapeutic tools to combat TB.

In *Mtb*, non-canonical DNA structures are implicated in the regulation of virulence genes, stress responses and drug resistance mechanisms. These structures form in regions of the genome that regulate essential cellular functions, making them prime candidates for therapeutic targeting. Several Non-B-DNA (Non Canonical) structures have been discovered, of these G-quadruplexes (GQ) and i-Motif (iM) sequences have drawn significant interest owing to their structural conformations and regulatory roles [3]. G4–DNA forming nucleic acid sequences have been discovered ubiquitously in both eukaryotes and prokaryotes. This suggests that GQ structures may be essential for regulating a range of cellular functions, such as transcription, translation and replication. Further it is suggested that the genomes of slow growing pathogenic species like *Mtb* are especially enriched for G4 sequences[4]. As a result, GQ-DNA structures are acknowledged as potential drug binding targets.

Higher order plants contain flavonoids, which are polyphenolic secondary metabolites. Flavonols are the most prevalent type of flavonoid of the many subclasses present in plants [5]. Owing to this, the human diet naturally contains significant levels of flavonoids that can be found in many fruits and vegetables [5]. They exhibit high pharmacological potency and low cytotoxicity making them viable alternatives to synthetic therapeutic drugs which are often associated with a high degree of cytotoxicity [6]. This makes them ideal therapeutic drugs with naturally high bioavalibity [7]. Both Quercetin (Que) and Kaempferol (Kae) have been shown to synergistically enhance the effects of conventional TB drugs while reducing drug associated cytotoxic effects however the bimolecular mechanisms remain unidentified [8,9].

Drug design must therefore focus on ensuring preferential recognition of these specific non canonical DNA structures, differentiating between GQ/iM and duplex DNA to prevent off target effects. Given their great potential for targeted therapeutic treatments at the genomic level, the search for G4/iM binding ligands is crucial. There is a growing body of evidence that supports the role of G4/iM DNA in regulating key cellular functions reinforcing the potential of phytochemicals as a viable therapeutic strategy. Building on this, the research into targeting GQ/iM DNA structures in *Mycobacterium tuberculosis* (*Mtb*) presents a considerable potential for combating TB-related drug resistance and virulence. In *Mtb*, non-canonical DNA structures are involved in the regulation of virulence genes, stress responses, and mechanisms of drug resistance. Additionally these structures occur in genomic regions within *Mtb* that control essential cellular functions, making them ideal candidates for therapeutic intervention. Identifying compounds that bind as well as stabilize these structures would result in replicative and transcriptional down regulation of key genes within Mtb disrupting cellular functions. Investigating phytocompounds as G4 and iM ligands could lead to the discovery of new, innovative therapeutics that could potentiate the effects of conventional anti-TB therapies.

In the present investigation, we explore the interaction between phytomolecules and non-canonical DNA structures by biophysical approaches such as UV-Vis absorption studies, binding constant determination and UV/thermal melting studies.

## Methods

### Identification of Non-Canonical Structures within *Mtb*

G Quadruplex and i-Motif DNA sequences within key TB genes were identified from available M.tb WGS (NC_000962.30) (CP023628.1 MDR) obtained from NCBI using several online tools such as iMseeker, QGRS and PQS Finder. The DNA sequences that have a high tendency to form non-canonical structures are selected. GQ sequences with (G_≥3_ N_1-7_)^4^ were selected along with a Human duplex DNA sequence as control to assess specificity towards *Mtb* sequences.

Additionally, The Comprehensive Resistance Prediction for Tuberculosis: an International Consortium (CRyPTIC) repository was used to select various drug resistant and drug susseptible strains that are prevalent within clinical cases in India that were sequenced in house (at The Foundation for Medical Research) listed in **Table 1**. The selected non canonical sequences were aligned using NCBI BLASTn with these strains to check for conservation. This would ensure that the selected sequences remain stable and viable therapeutic targets irrespective of strain and resistance profiles.

**Table 1.** Strain type and resistance profile of prevalent Indian *Mtb* strains obtained from the CRyPTIC repository screened for homology with selected G4/iM sequences. (Lineages: L1 – Indo Oceanic, L2-East Asian, L3 – East African Indian, L4-Euro-American Lineage.)

| **Resistance Profile** | **Strain Id.** | **Lineage** |
| --- | --- | --- |
| Drug Sensitive | 628076 | 1.1.2 |
| Drug Sensitive | 631135 | 3 |
| Drug Sensitive | 631903 | 1.1.2 |
| Drug Sensitive | 632518 | 3 |
| Drug Sensitive | 632635 | 1.2.2.2 |
| Drug Sensitive | 632664 | 3 |
| Drug Sensitive | 633210 | 1.1.2 |
| Drug Sensitive | 633418 | 4.2.2.2 |
| Drug Sensitive | 633681 | 1.1.2 |
| Drug Sensitive | 714176 | 3 |
| Drug Resistant (Pre XDR) | 630821 | 2.2.1 |
| Drug Resistant (Pre XDR) | 631235 | 2.2.1.1 |
| Drug Resistant (Pre XDR) | 631450 | 2.2.1 |
| Drug Resistant (Pre XDR) | 631621 | 2.2.1.1 |
| Drug Resistant (Pre XDR) | 631993 | 2.2.1 |
| Drug Resistant (Pre XDR) | 632619 | 2.2.1 |
| Drug Resistant (Pre XDR) | 632901 | 3 |
| Drug Resistant (Pre XDR) | 633291 | 2.2.1 |
| Drug Resistant (Pre XDR) | 633497 | 2.2.1 |
| Drug Resistant (Pre XDR) | 633541 | 1.1.2 |

## Materials and DNA Preparation

*Mtb* G4 DNA sequences, iM sequences and the duplex DNA sequence were procured from Eurofins Scientifics. Quercetin (Q4951) and kaempferol (60010) were obtained from Sigma Aldrich. **(Table 2, Figure 1)**. The solvents used were of spectroscopic grade. Stock solutions of Quercetin and Kaempferol were prepared in methanol (because of low solubility in an aqueous medium), and the final experimental concentrations of all flavonoids were kept on the order of 10^−6^ M, methanol <1% (v/v). The desalted oligonucleotides were dissolved in double distilled water and stored at 4 °C. G4/iM DNA solutions were prepared by taking a requisite amount of DNA from the main stock to the buffer solution, and the resulting solutions were annealed by heating at 95 °C for 5 min. The solutions were then slowly cooled to room temperature and equilibrated overnight at 4 °C to allow for sequence folding. All experiments were carried out using Sequence specific GQ or iM Buffers. GQ Buffer composition being 50 mM KCl, 10 mM KH_2_PO4, and 1 mM K_2_EDTA (pH 7.4, 25 °C) and iM Buffer comprising of 50 mM KCl, 10 mM KH_2_PO_4_, and 1 mM K_2_EDTA (pH-5.4, 25 °C).

**Img 1:**
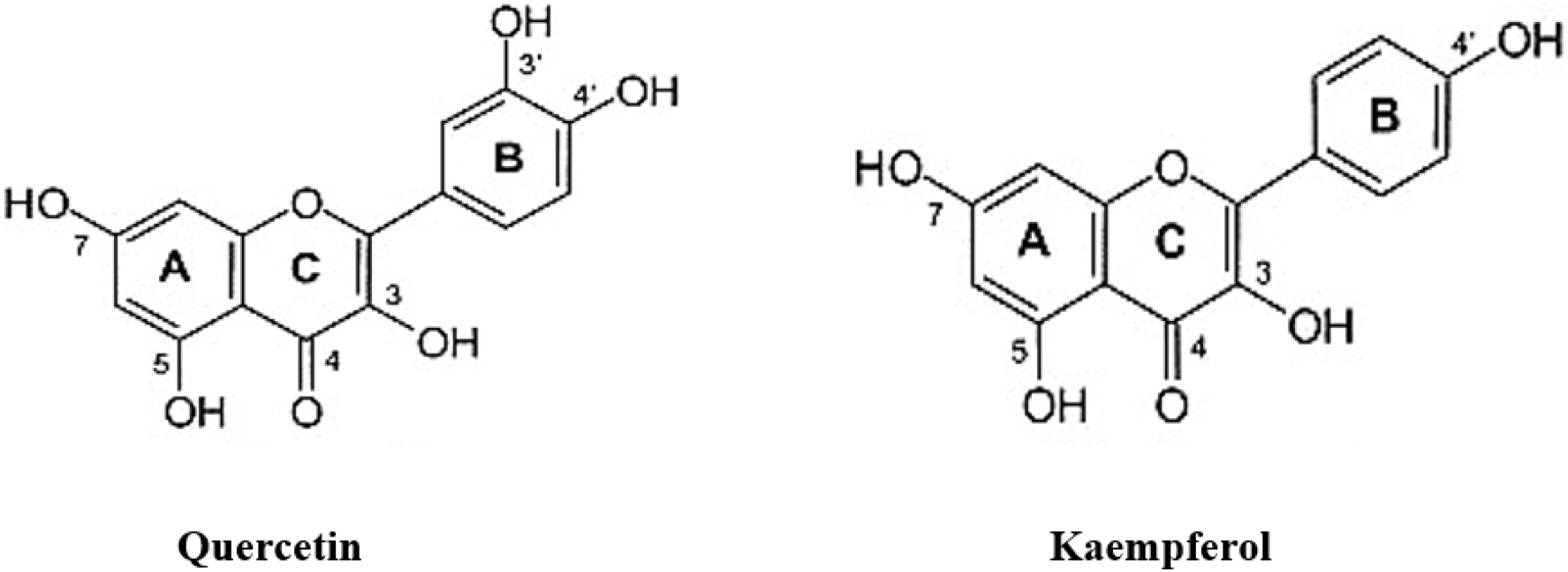
Schematic representation of structures of Quercetin and kaempferol.

**Table 2:**
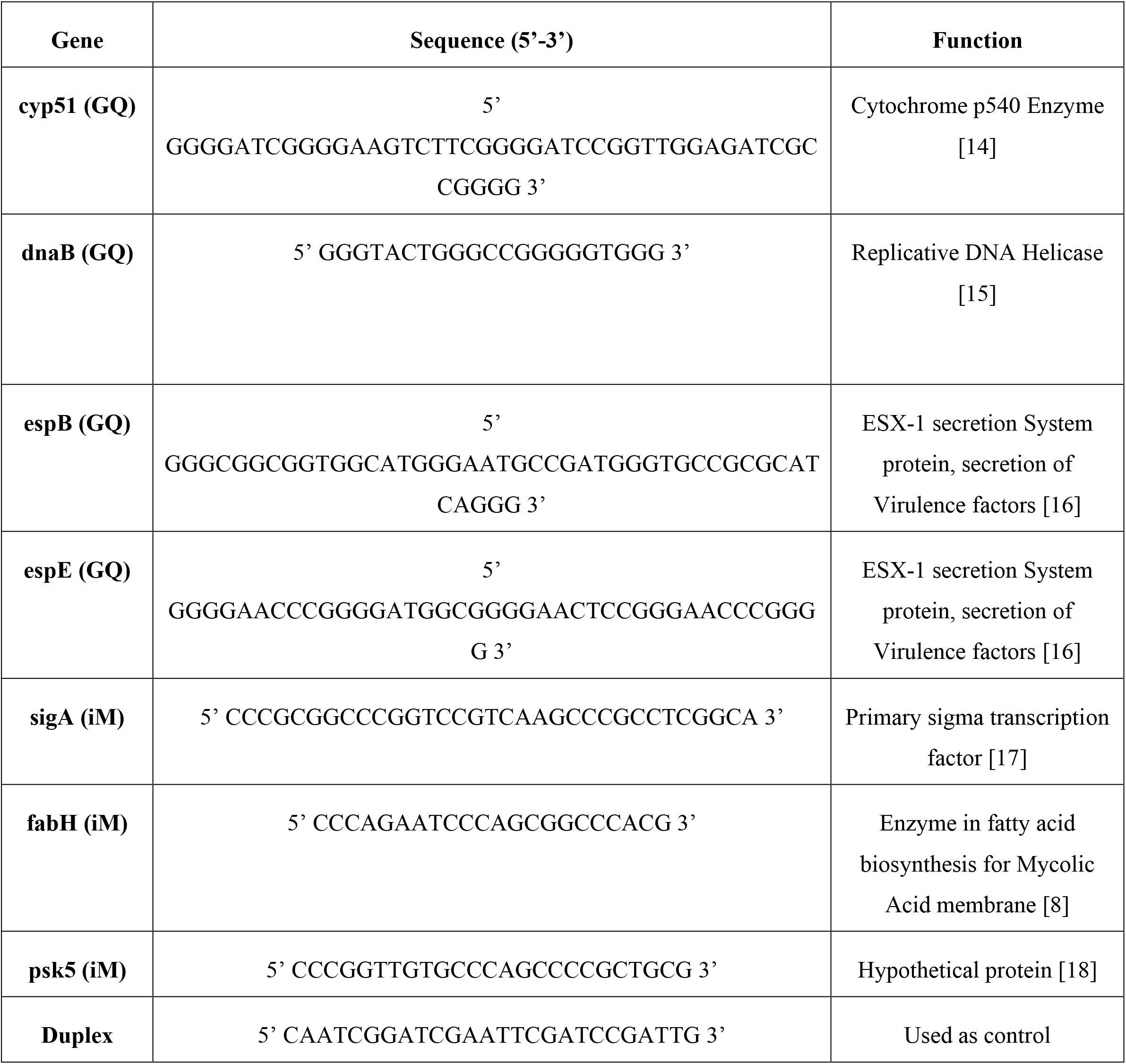
DNA sequences used in this study. Identification of conserved GQ and iM forming sequences within *Mtb*.

**Figure 1.**
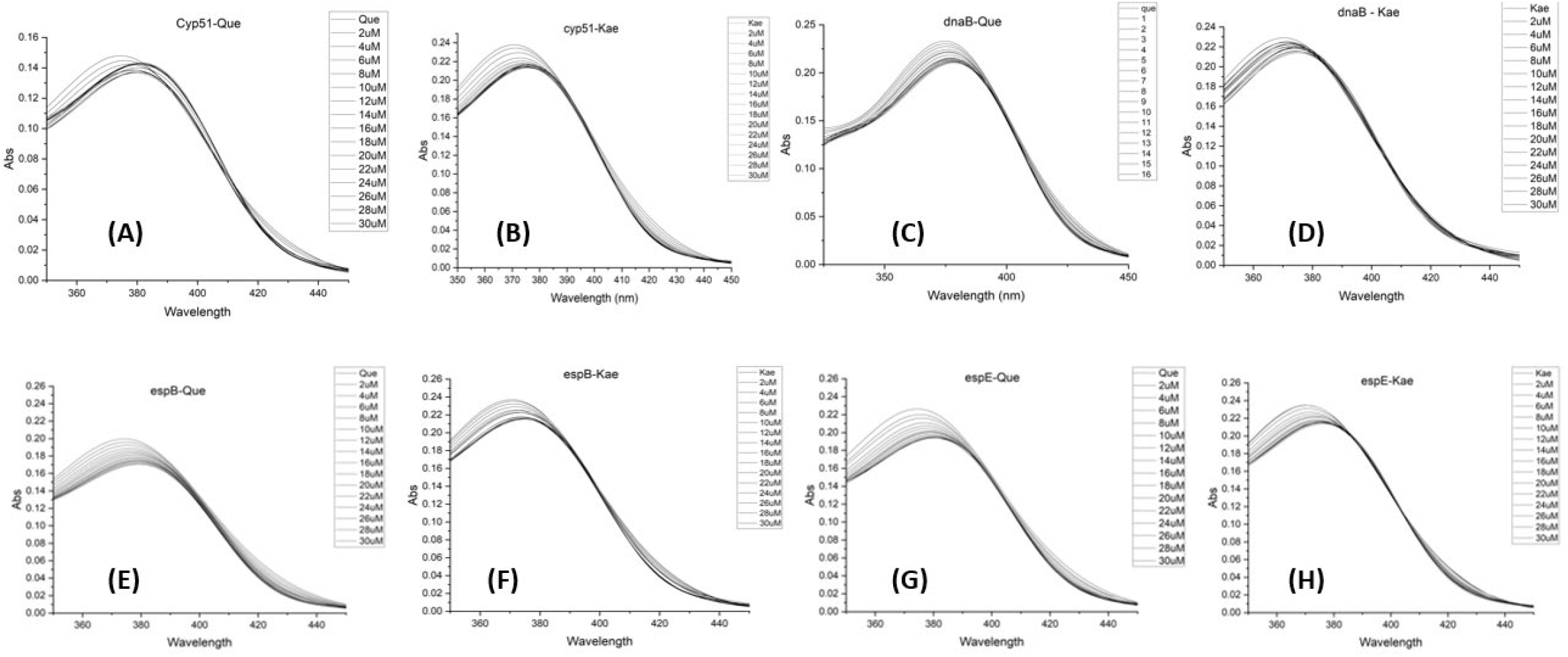
UV-Vis absorption spectra of Que and Kae (15 μM) in the absence and presence of successive additions of cyp51 GQ, dnaB GQ, espB GQ and espE GQ (0-30 μM) [1A-1H].

### UV−Vis Absorption Studies

UV−Vis absorption spectra were recorded with a Jasco V-630 spectrophotometer. Measurements were performed in matched quartz cuvettes of 1 cm path length. For both experiments the ligand (Que or Kae) concentrations were kept constant (15 μM) and an increasing DNA concentration was titrated (2 μM - 30 μM). The UV−Vis absorption spectra of the flavonoid Que and Kae were examined in the presence of GQ and iM DNA constructs in order to acquire a preliminary understanding of the binding associations [8]. Que and Kae follow a conventional diphenyl propane framework, which is the basis for their chemical compositions, made up of two benzene rings. The rings, A (benzoyl moiety) and B (cinnamoyl moiety), connected by a heterocyclic pyran or pyrone (with a double bond) ring (C) in the middle. This core structure may also contain a number of hydroxyl groups (OH) and/or additional substituents. Several recent reports suggest that the position and number of OH groups on the B–ring of flavonoids are important for their DNA, RNA, and protein binding properties. Both flavonols show structural similarities, however Quercetin has an additional hydroxyl (−OH) group at the 3’ position in its B ring, possibly enhancing electron donation and radical scavenging via resonance delocalization. In general, the absorption profiles of the flavonoid have revealed two bands of absorption (band I and band II) with Band I displaying the light absorption of the cinnamoyl moiety (B+C ring), while band II reflects the absorbing capability of the benzoyl moiety (A+C ring) [9].

### Binding Constant Determination

Determining the binding constant between a phytochemical and non-canonical DNA structures is essential for evaluating the strength and specificity of their interaction, which influences the drug’s effectiveness, stability, and selectivity. The binding constant (K_b_) quantifies the affinity between the phytochemical and a specific DNA structure, with a higher constant indicating a stronger interaction with the target sequences suggesting strong potential as therapeutic drugs. Measuring the binding constant helps assess whether the phytochemical selectively interacts with these structures instead of typical B-DNA. Understanding the binding affinity can provide insights into whether the interaction is intercalative, groove-binding, or external stacking. This information is crucial for predicting the potential biological effects of the phytochemical. To calculate the binding constant (K_b_) of flavonoids (quercetin and kaempferol) - GQ/iM DNA interaction, we used the following equation based on absorbance variation [10]

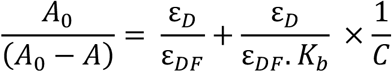

Where A_0_ and A represent the absorbance of free and Que/Kae-treated GQ/iM DNA and εD and εDF are the molar extinction coefficients of GQ/iM DNA and GQ/iM DNA-flavonoid complex, respectively, and C is the concentration of flavonoids. The double reciprocal plot of 1/(A−A_0_) vs 1/C was linear, and the binding constant (K_b_) could potentially be calculated using the intercept-to-slope ratio.

### Thermal Melting Studies

Thermal denaturation experiments provided an alternative method to evaluate the interaction between Que, Kae and experimental cyp51 GQ, dnaB GQ, SigA iM and duplex DNA structures. At specific temperatures, localized disruptions in the DNA helix occur, causing the strands to separate [11], [12], [13]. The presence of small molecules like flavonoids has been shown to raise the melting temperature (T_m_) of GQ/iM DNA by strengthening the helical structure through base stacking, as binding through grooves or electrostatic interactions does not significantly affect the melting temperature. Thermal denaturation experiments were conducted using a Hitachi-UH5300 spectrophotometer, equipped with a Julabo F12 water stirrer and a 3J1-0104 water pumping cell holder apparatus, employing a 1-centimeter path length quartz cuvette. The samples were heated at a constant rate of 1°C per minute, from 30°C to 98°C. The absorbance values of μM GQ and iM DNA, both alone and in the presence of 20 μM quercetin and 10 μM kaempferol, were monitored at 295 nm as the temperature increased from 30°C to 98°C, with measurements taken every 0.5°C. The experiments were performed in the respective GQ and iM buffer solutions described earlier. The UV melting curves were normalized and analyzed using the Origin Pro 8 software.

## Results

### Identification of Non-Canonical Structures within *Mtb*

GQ and iM sequences within the *Mtb* genome were identified. These were narrowed down to seven sequences with key roles in *Mtb* regulation and resistance listed in **Table 2**. These sequences were also found to be highly conserved within prevalent TB strains in India identified from the CRyPTIC repository. They were identically conserved in drug resistant and drug susceptible strains across all 4 lineages listed in **Table 1**.

### UV−Vis Absorption Measurements

The decreased absorption (hypochromicity), coupled with a red shift (bathochromicity) in the absorption maxima, showed a possible polarity transfer around the flavonoid compounds as a consequence of their binding relationship with DNA structures. In the presence of increasing concentrations of cyp51 GQ, dnaB GQ, espB GQ, espE GQ, SigA iM, fabH iM, psk5 iM DNA, the absorption spectra of Que and Kae displayed a significant decrease in absorbance, along with a noticeable red shifting in the wavelength of the absorbance maxima. The decreased absorption together with the prominent red shift at the absorption maxima of Que and Kae, demonstrated that the binding interaction with cyp51 GQ, dnaB GQ, espB GQ, espE GQ, SigA iM, fabH iM, psk5 iM DNA caused polarity swapping around the Que and Kae molecules. These findings also revealed that Que and Kae detect a hydrophobic atmosphere within these DNA matrices, where it is sheltered from the polar aqueous environment. The discovered bathochromic consequences were most likely caused by the interactions of the unoccupied antibonding orbital (π*) of Que and Kae with the bonding orbital (π) of the DNA bases, resulting in a π−π* conjugative pairing, a decrease in the π−π* transition dynamism, and a red shift in the adsorption profile. The unoccupied π*-orbital of Que and Kae, on the other hand, has been partially filled with electrons, which are decreasing the transition probability and causing hypochromism because the lower the probability, the lesser the molar absorption proportion.

These data clearly demonstrated the potential for Que and Kae to be associated with cyp51 GQ, dnaB GQ, espB GQ, espE GQ, SigA iM, fabH iM, psk5 iM DNA compared to other experimental DNAs and duplex DNA. Based on the magnitude of the spectrum modifications in the absorption profile in our experimental findings, we infer that Que and Kae interact with the DNA through a strong stacking mode of binding owing to the significant hypochromic shifts along with significant red shifting in the UV−Vis absorption profile.

Based on the UV-Vis absorption studies, we determined that Que and Kae preferentially interacts with cyp51 GQ, dnaB GQ, espB GQ, espE GQ, SigA iM, fabH iM, psk5 iM DNA over other experimental GQ, iM and duplex DNA’s **(Table 3, Figures 1**,**2)**.

**Table 3.** Spectral parameters of interactions obtained from UV Visible Absorption experiments.

| **Sequence** | **Ligand** | **Hypochromicity (%)** | **Red Shift (nm)** |
| --- | --- | --- | --- |
| **Duplex** | **Kae** | **5.69** | **0.9** |
|  | **Que** | **12.32** | **1.4** |
| **cyp51 (GQ)** | **Kae** | **9.28** | **6.1** |
|  | **Que** | **2.72** | **7.3** |
| **dnaB (GQ)** | **Kae** | **6.55** | **4.7** |
|  | **Que** | **9.48** | **3.1** |
| **espB (GQ)** | **Kae** | **9.70** | **6** |
|  | **Que** | **14.57** | **6.2** |
| **espE (GQ)** | **Kae** | **8.12** | **6.6** |
|  | **Que** | **14.16** | **6.8** |
| **sigA (iM)** | **Kae** | **17.12** | **6.9** |
|  | **Que** | **18.37** | **8.7** |
| **fabH (iM)** | **Kae** | **18.50** | **8** |
|  | **Que** | **29.27** | **7.2** |
| **psk5 (iM)** | **Kae** | **20.57** | **16** |
|  | **Que** | **22.18** | **11.4** |

**Figure 2.**
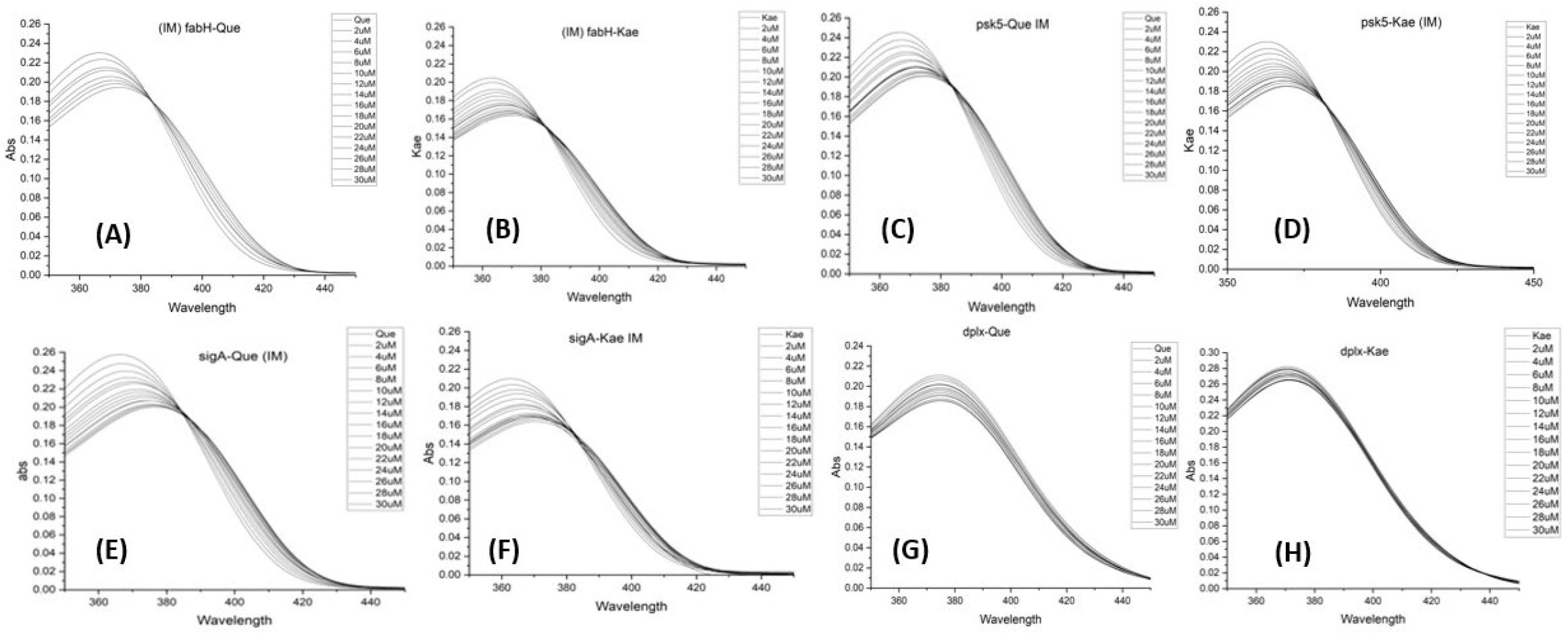
UV-Vis absorption spectra of Que and Kae (15 μM) in the absence and presence of successive additions of fabH iM, psk5 iM, SigA iM and duplex DNA (0-30 μM) [2A-2H].

### Binding Constant determination

In this study, we calculated the binding constant (K_b_) values for a series of flavonoid (Que/Kae)-GQ/iM DNA complexes by analysing the variations in DNA absorbance as a function of flavonoid concentration. The calculated binding constants are indicative of the strength and specificity of the interactions between the flavonoids and the different DNA sequences, which are crucial for understanding the mode of action of these flavonoids. The binding constants (K_b_) were derived from the changes in absorbance, reflecting how the flavonoids interact with the DNA in solution. These binding constants are essential for comparing the relative affinities of the flavonoids (Que and Kae) for various DNA targets, including cyp51 GQ, dnaB GQ, espB GQ, espE GQ, SigA iM, fabH iM, and psk5 iM DNA. The calculated binding constants are tabulated in **Table 4 and Figures 3A-3G and 4A-4G**.

**Table 4.**
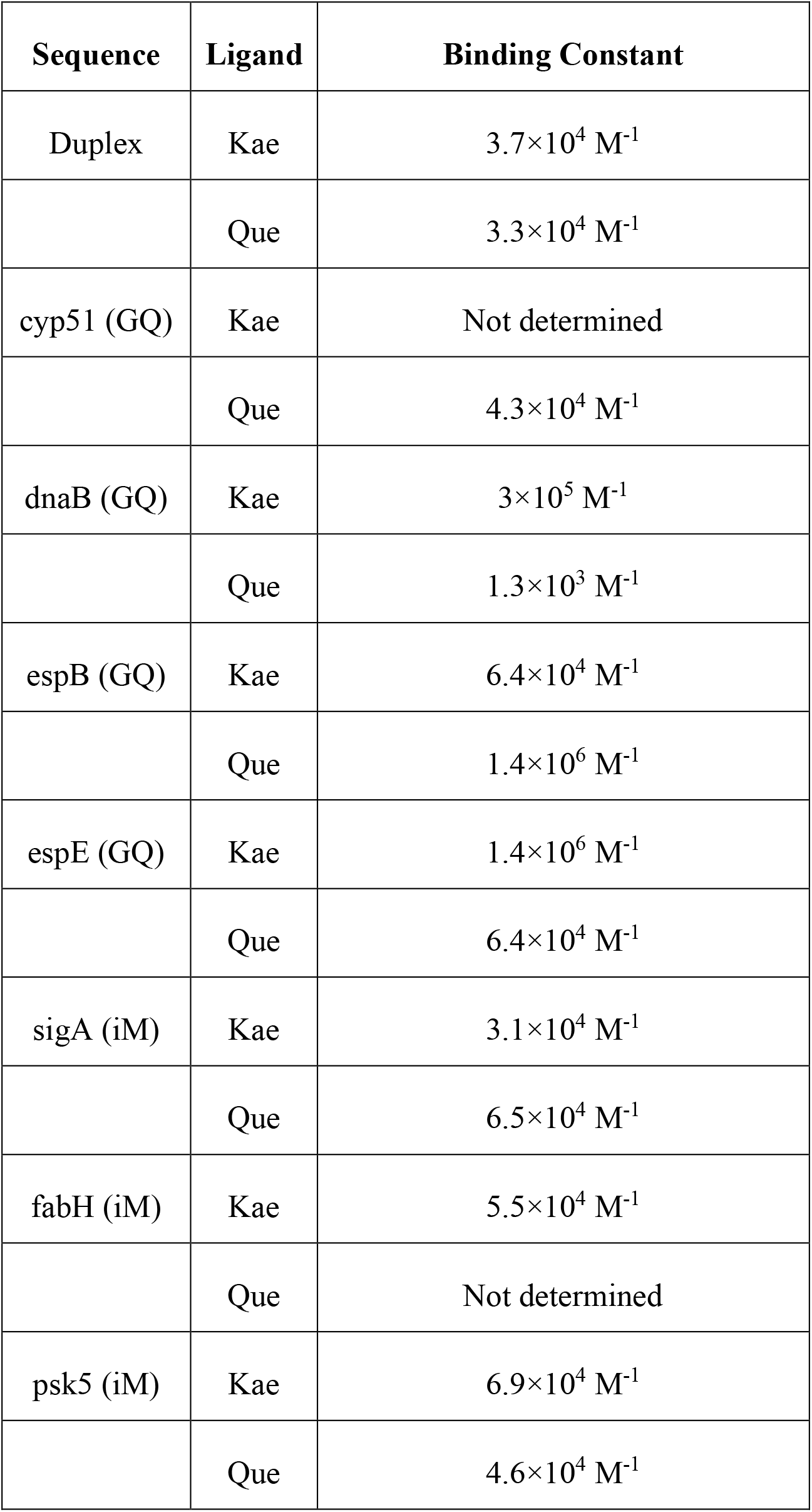
Binding constant determination from UV Visible Absorption experiments.

**Figure 3.**
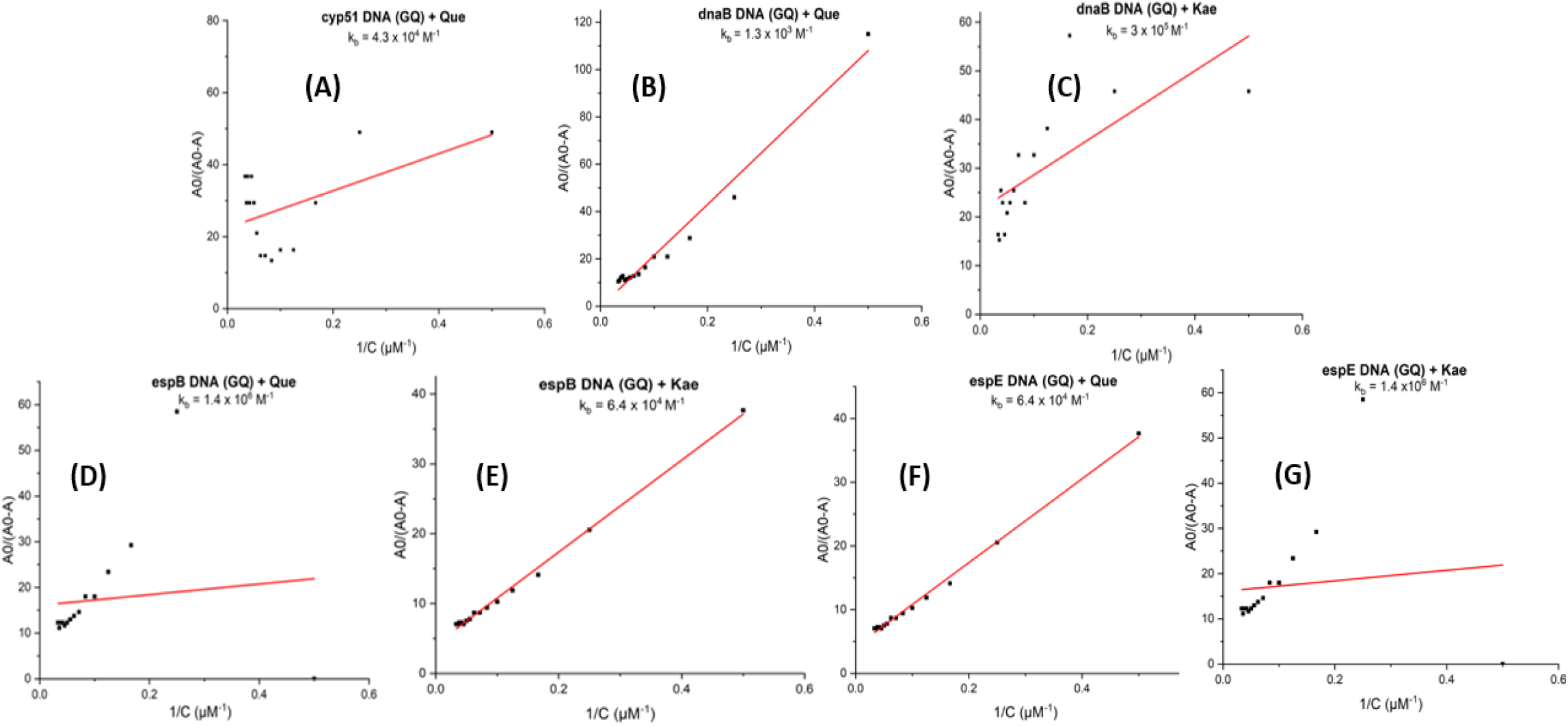
Double reciprocal plot for binding constant determination of cyp51 GQ, dnaB GQ, espB GQ and espE GQ DNA in the presence of Que and Kae (4A-4G).

**Figure 4.**
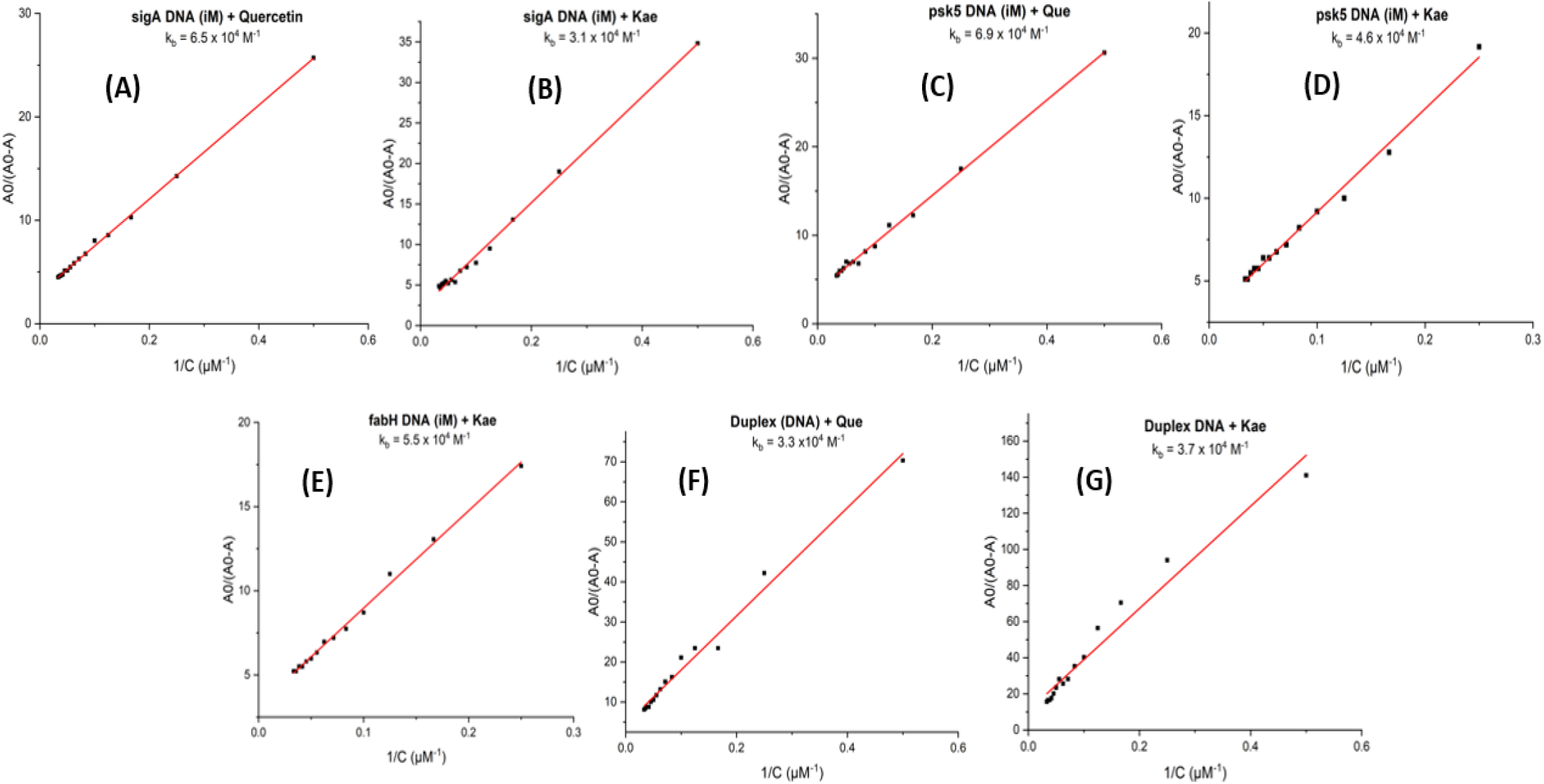
Double reciprocal plot for binding constant determination of SigA iM, psk5 iM, fabH iM and duplex DNA in the presence of Que and Kae (4A-4G).

A strong interaction of the phytocompounds with the selected non-canonical DNA structures is observed. In conclusion, the binding constants calculated in this study are indicative of the moderate (10^3 –^ 10^4^ M^-1^) and strong (10^5 –^ 10^7^ M^-1^) binding and stabilizing interactions between Que and Kae and GQ/iM DNA, shedding light on how these compounds selectively target specific GQ/iM DNA structures [19].

Going further cyp51 (GQ), dnaB (GQ) and sigA (iM) sequences were used for thermal melting studies owing to their increased specificity for the tested phytocompounds and significant in vivo functions.

### Thermal Melting Studies

Thermal melting (T_m)_ experiments explore how Que and Kae impacts the stability of cyp51 GQ, dnaB GQ and SigA iM. The T_m_ for each transition was calculated by identifying the midpoint of the melting curve. The absorption at 295 nm of cyp51 GQ, dnaB GQ and SigA iM bases was used to determine the T_m_. **Figures 5A-5H** display melting curves and temperature changes with and without Que and Kae. The addition of 20 μM Que (2:1, Ligand:DNA concentration)notably increased the thermal stability of cyp51 GQ, dnaB GQ and SigA iM DNA by 7.6, 7.9 and 7 °C **(Figure 5A, 5C, 5F)**. The addition of 10 μM Kae (1:1, Ligand: DNA concentration) also increased the thermal stability of cyp51 GQ, dnaB GQ and SigA iM DNA by 8.6, 5.5 and 8.3 °C **(Figure 5B, 5D, 5E)**. However, the addition of 20 μM Que and 10 μM Kae have no impact on the thermal stability of duplex DNA (**Figure 5G, 5H, Used as control)**.

**Figure 5.**
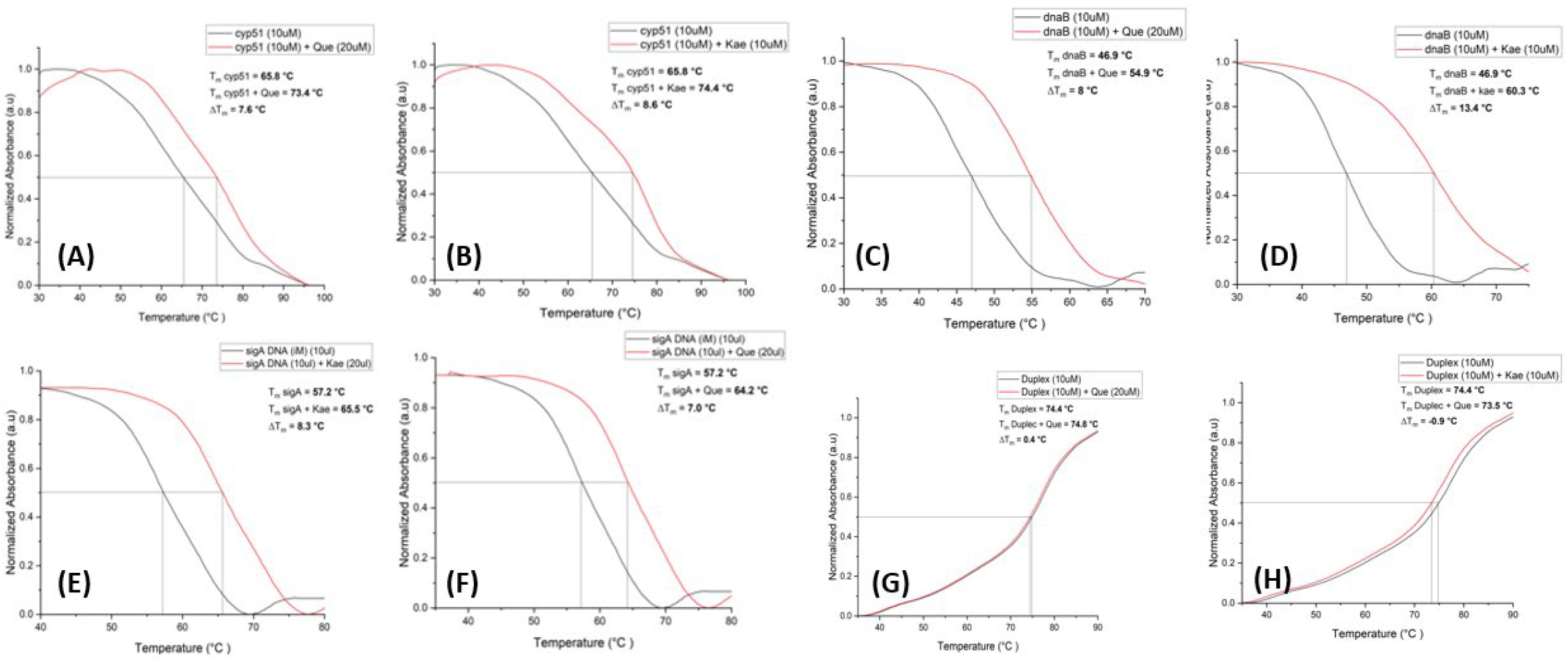
Thermal melting profiles of cyp51 GQ, dnaB GQ, SigA iM and duplex DNA (10 μM) alone and in the presence of Que (10 μM) and Kae (20 μM). Variations of the GQ/iM DNA melting temperature (ΔT_m_) in °C induced by Que/Kae.

These results eliminated the possibility of Que and Kae binding through a loop or groove (exterior binding mode) and provided stronger evidence for a robust stacking and stabilizing interactions with cyp51 GQ, dnaB GQ and SigA iM DNA.

## Discussion

Non canonical DNA structures have been investigated for the evolutionary functions and roles in diseases since their discovery[20], [21] [22]. The study of such structures within the human genome especially in regards to their roles in cancer has increasing gained traction [22]. However the role of these structures in prokaryotes and pathogenic micro-organisms has until recently remained under explored.

This study undrescores the potential of Quercetin and Kaempferol as sequence and pathogen specific ligands, showing preferential binding to the identified G-quadruplex (GQ) and i-motif (iM) structures, particularly within *Mycobacterium tuberculosis* (Mtb). This investigation revealed that quercetin (Que) and kaempferol (Kae) selectively bind to the cyp51, dnaB, espB, espE GQ as well as sigA, fabH and psk5 iM DNA sequences, showing a stronger affinity compared to other tested GQ, iM, and duplex DNA sequences. The interactions between these phytochemicals and *Mtb* DNA structures reveal their significant binding affinity depicted by the changes in UV-Vis absorption spectra and supported by increased thermal stability of these DNAs. These findings suggest that flavonoids not only bind preferentially to these GQ and iM structures but also stabilize these non-canonical forms, potentially interfering with critical cellular processes such as gene regulation and resistance mechanisms. The ability of these flavonoids to enhance the effects of conventional anti-TB drugs, while minimizing cytotoxicity, presents a novel therapeutic avenue to overcome the challenges posed by drug-resistant strains of *Mtb*[20]. When combined in a cocktail, Que and Kae may act synergistically, enhancing their affinity for GQ/iM DNA structures across the TB genome. By targeting these structures, the flavonoids may downregulate the expression of critical resistance genes or interfere with DNA replication and repair mechanisms, thus reducing the pathogen’s ability to develop resistance to existing TB drugs. Overall, this research contributes to the growing body of evidence supporting the use of natural compounds for targeting key genomic structures in drug-resistant pathogens, paving the way for more effective and safer treatments for tuberculosis [4], [22].

## Acknowledgements

The study was supported by Core Funds of the Foundation for Medical Research, Mumbai. We acknowledge Mr. Atanu Manna, University of Calcutta for his initial bioinformatics inputs in the study.

